# Receptor utilization of angiotensin converting enzyme 2 (ACE2) indicates a narrower host range of SARS-CoV-2 than that of SARS-CoV

**DOI:** 10.1101/2020.06.13.149930

**Authors:** Ye Qiu, Qiong Wang, Jin-Yan Li, Ce-Heng Liao, Zhi-Jian Zhou, Xing-Yi Ge

## Abstract

Coronavirus pandemics have become a huge threat to the public health worldwide in the recent decades. Typically, SARS-CoV caused SARS pandemic in 2003 and SARS-CoV-2 caused the COVID-19 pandemic recently. Both viruses have been reported to originate from bats. Thus, direct or indirect interspecies transmission from bats to humans is required for the viruses to cause pandemics. Receptor utilization is a key factor determining the host range of viruses which is critical to the interspecies transmission. Angiotensin converting enzyme 2 (ACE2) is the receptor of both SARS-CoV and SARS-CoV-2, but only ACE2s of certain animals can be utilized by the viruses. Here, we employed pseudovirus cell-entry assay to evaluate the receptor-utilizing capability of ACE2s of 20 animals by the two viruses and found that SARS-CoV-2 utilized less ACE2s than SARS-CoV, indicating a narrower host range of SARS-CoV-2. Meanwhile, pangolin CoV, another SARS-related coronavirus highly homologous to SARS-CoV-2 in its genome, yet showed similar ACE2 utilization profile with SARS-CoV rather than SARS-CoV-2. To clarify the mechanism underlying the receptor utilization, we compared the amino acid sequences of the 20 ACE2s and found 5 amino acid residues potentially critical for ACE2 utilization, including the N-terminal 20^th^ and 42^nd^ amino acids that may determine the different receptor utilization of SARS-CoV, SARS-CoV-2 and pangolin CoV. Our studies promote the understanding of receptor utilization of pandemic coronaviruses, potentially contributing to the virus tracing, intermediate host screening and epidemic prevention for pathogenic coronaviruses.

## Introduction

Coronaviruses are enveloped non-segmented positive sense RNA viruses. In the recent two decades, human coronaviruses (HCoVs) have caused to at least three major pandemics and pose a huge threat to the public health worldwide (1). Until now, seven different HCoVs have been identified and all of them belong to the family *Coronaviridae* of the order *Nidovirales*. Especially, the species *Severe acute respiratory syndrome-related coronavirus* (SARSr-CoV) of the genus *Betacoronavirus* contains the most pathogenic HCoVs, including severe acute respiratory syndrome coronavirus (SARS-CoV) and severe acute respiratory syndrome coronavirus 2 (SARS-CoV-2) (2). SARS-CoV caused the SARS pandemic in years of 2002 – 2003, resulting in more than 8,000 clinical cases with a mortality of 10% (3, 4). After the SARS pandemic, plenty of research and measurements have been done to prevent the emergence and re-emergence of coronavirus epidemics, including transmission tracing, pathogenesis studies and therapy development (5). Nevertheless, 17 years later, a novel human coronavirus, SARS-CoV-2, brought a much more severe and widespread pandemic of Coronavirus Disease 2019 (COVID-19) to the world and this pandemic is lasting until now (6, 7). Up to May 25, 2025, about 5 month after the first reported case of COVID-19, SARS-CoV-2 has caused 5,432,302 confirmed infections and 342,318 deaths worldwide, far surpassing the SARS pandemic. So far, most countries around the world are suffering from the public health crisis, medical system overload and economic loss due to the pandemic. Among common pathogenic viruses, SARS-CoV-2 emerges a high transmissibility whose R0 value is currently estimated as 2.3 but could be as high as 5.7 when more infection cases are identified (8). The high transmissibility of SARS-CoV-2 is probably a major reason for the rapid development of COVID-19 epidemic. In order to relieve the current pandemic and prevent HCoV epidemics in the future, more studies on the transmission of pathogenic HCoV is urgently required.

Interspecies transmission from wide animals to humans is a major cause leading to the epidemics of highly pathogenic coronaviruses, such as SARS-CoV and SARS-CoV-2. Previous virus tracing studies have shown that Chinese horseshoe bats are natural reservoirs of SARS-CoV (9, 10), and a recent phylogenetic analysis has revealed that SARS-CoV-2 might also be originated from bat-SARSr-CoV (11). Some small mammals, such as civets (*Paguma larvata*) and raccoon dogs (*Nyctereutes procyonoides*), can serve as the intermediate hosts of SARS-CoV and might be he direct sources of the SARS epidemic in early 2003 due to the successful isolation of SARS-CoV from these animals (12). Similarly, SARS-CoV-2-like CoVs were detected in Malayan pangolins, indicating pangolins might serve as an intermediate host for SARS-CoV-2 (13, 14). The intermediate host animals usually play critical roles in the interspecies transmission from natural virus reservoirs to humans and community transmission of viruses. The host range of a virus is an essential factor determining its intermediate hosts. Thus, research on the host range of viruses is of great importance for virus tracing and epidemic control.

A main factor determining the host range of viruses is the recognition and binding between viral particles and their receptors on the host cells. Angiotensin converting enzyme 2 (ACE2) is utilized by SARS-CoV and SARS-CoV-2 as their cellular receptor (1, 15). Discovered in the year of 2000, ACE2 was initially identified as an exopeptidase expressed in vascular endothelial cells in the heart and the kidney that catalyses the conversion of angiotensins (16, 17). ACE2 is highly conserved and ubiquitously expressed in most vertebrates, but not all ACE2s can serve as the receptor for SARS-CoV and SARS-CoV-2, which is a key determinant of the host range of these two viruses. For instance, SARS-CoV can use mouse ACE2 as its receptor but SARS-CoV-2 cannot, indicating that mouse is a potential host for SARS-CoV but not for SARS-CoV-2 (1). Our previous study predicted the ACE2 utilization of SARS-CoV-2 and showed that SARS-CoV-2 could potentially utilize most mammalian ACE2s except murine ACE2s and even some avian ACE2s. Meanwhile, we also predicted 9 amino acid (aa) residues in ACE2 critical for SARS-CoV-2 utilization (18). However, this study was mainly based on the aa analysis and lacked experimental evidence, which could be hardly referred to clarify the host range of SARS-CoV-2.

In this study, we ectopically expressed ACE2 of 20 different animals in HeLa cells, a cell line lacking of ACE2 expression naturally, and then infected the cells with HIV-based pseudoviral particles carrying coronavirus spike proteins to test their utilization of these ACE2s. The result showed that both SARS-CoV and SARS-CoV-2 could use most mammalian ACE2s as their receptors but not fish or reptilian ACE2s. Interestingly, similar to mouse ACE2, SARS-CoV but not SARS-CoV-2 was capable of using chicken ACE2, indicating a narrower host range of SARS-CoV-2, especially in murine and birds. We also tested pangolin CoV pseudovirus with spike highly similar to SARS-CoV-2. To our surprise, pangolin CoV showed an ACE2 utilization profile similar to SARS-CoV rather than SARS-CoV-2. By alignment of the aa sequence of the 20 ACE2 orthologs, we further confirmed several aa residues (among the 9 residues predicted previously) critical for SARS-CoV-2 utilization, including T20, K31, Q42 and Y83, and excluded some critical aa residues, such as Y41 and K68. Especially, T20 of ACE2 probably played critical roles in spike-ACE2 binding by interacting with S477 and T478 within the receptor-binding motif (RBM) of SARS-CoV-2 spike protein, but it was not necessary for the binding of ACE2s with SARS-CoV and pangolin CoV spikes. These aa residues might partially determine the unique receptor utilization and host range of SARS-CoV-2. In summary, our study provides a more clear view of ACE2 utilization by SARS-CoV-2, which may contribute to a better understanding about the virus-receptor interaction and the host range of SARS-CoV-2.

## Results

### Validation of pseudovirus preparation and ACE2 expression

In this study, we investigated the ACE2 utilization of three SARSr-CoVs, including SARS-CoV-BJ01, SARS-CoV-2 and Pangolin CoV. Pangolin CoV was tested since pangolins were suspected to be a potential intermediate host of SARS-CoV-2 (14). Notably, the RBM of Pangolin CoV spike is almost identical with SARS-CoV-2 spike except for the 498^th^ aa residue, but they are different from SARS-CoV spike on multiple aa sites (**Figure 1A**).

**Figure 1.**
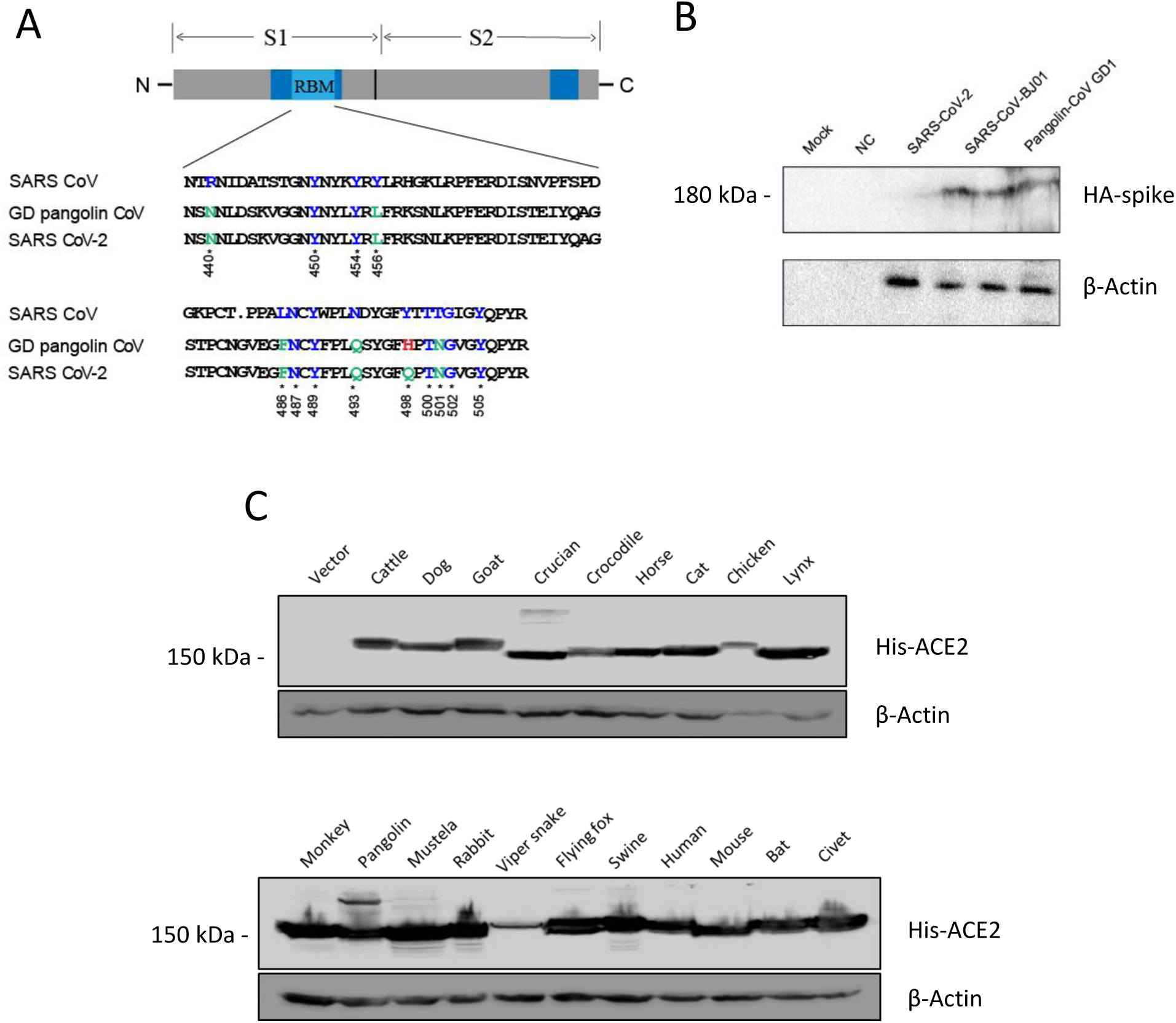
Validation of pseudovirues preparation and ACE2 expression. (A) Western blot Detection of spike proteins of SARS-CoV-BJ01, SARS-CoV-2 and Pangolin CoV in the package cells using an antibody against the HA tag conjugated to the viral spike proteins. β-actin was detected as the loading control. (B) Alignment of the amino acid sequences of the receptor binding motifs (RBM) of SARS-CoV, SARS CoV-2 and Pangolin CoV spike proteins. (C) Detection of different ACE2 orthologs in HeLa cells after transfecting the corresponding plasmids using an antibody against the 6XHis tag conjugated to the ACE2 proteins. β-actin was detected as the loading control.

In order to validate the successful preparation of the three pseudovirus, the HEK293T cells for virus package were lysed after virus harvest the cells were lysed and subjected to Western blot detection of HA-tagged spike proteins. As shown in **Figure 1B**, samples from all three package cells showed typical bands (about 180 kDa) of SARSr-CoV spike proteins, indicating the success of pseudovirus preparation.

Then, we continued to validate the ectopic expression of 20 ACE2s from different animals in HeLa cells, an ACE2-negative human cell line. We transfected the HeLa cells with plasmids harboring the coding gene of ACE2s from different animals. At 48 h post transfection, he cells were lysed and subjected to Western blot detection of 6*His-tagged ACE2 protein. As shown in **Figure 1C**, bands of all ACE2s (about 150 kDa) could be observed, indicating the successful expression of all ACE2s in HeLa cells.

### Difference in the ACE2 utilization by SARS-CoV and SARS-CoV-2

We transfected the HeLa cells with the 20 plasmids expressing different ACE2s individually or empty vector as a control. At 48 h post transfection, the cells were infected with SARS-CoV-BJ01, SARS-CoV-2 or Pangolin CoV pseudovirus. After 48 h of infection, the cells were lysed and subjected to luciferase assay to evaluate the cell-entry efficiency of the pseudoviruses mediated by different ACE2s. As shown in **Figure 2**, little luminescence signals could be observed in samples from HeLa cells transfected with empty vector and infected by any of the three pseudoviruses, indicating that native HeLa without ACE2 could not mediate the pseudovirus entry. Luminescence signals from cells expressing crucian, crocodile or viper snake ACE2 were also low, indicating that fish and reptilia ACE2s could barely mediate the pseudovirus entry. Cells expressing chicken or mouse ACE2 showed a strong luminescence signal when infected by the SARS-CoV-BJ01 pseudovirus but not by the SARS-CoV-2 pseudovirus, indicating that SARS-CoV-BJ01 could use chicken or mouse ACE2 for cell entry but SARS-CoV-2 could not. Pangolin CoV was capable of utilizing both chicken and mouse ACE2s, but its utilizing efficiency of chicken ACE2 was much lower than SARS-CoV-BJ01. On the contrary, SARS-CoV-2 pseudovirus ignited a stronger luminescence than SARS-CoV-BJ01 or Pangolin CoV pseudovirus in cells expressing bat ACE2, indicating the highest utilizing capability of bat ACE2 by SARS-CoV-2. For the other ACE2s, infection of all the three pseudoviruses led to strong luminescence signals, implying that all the three SARSr-CoV were capable of utilizing a broad range of ACE2s.

**Figure 2.**
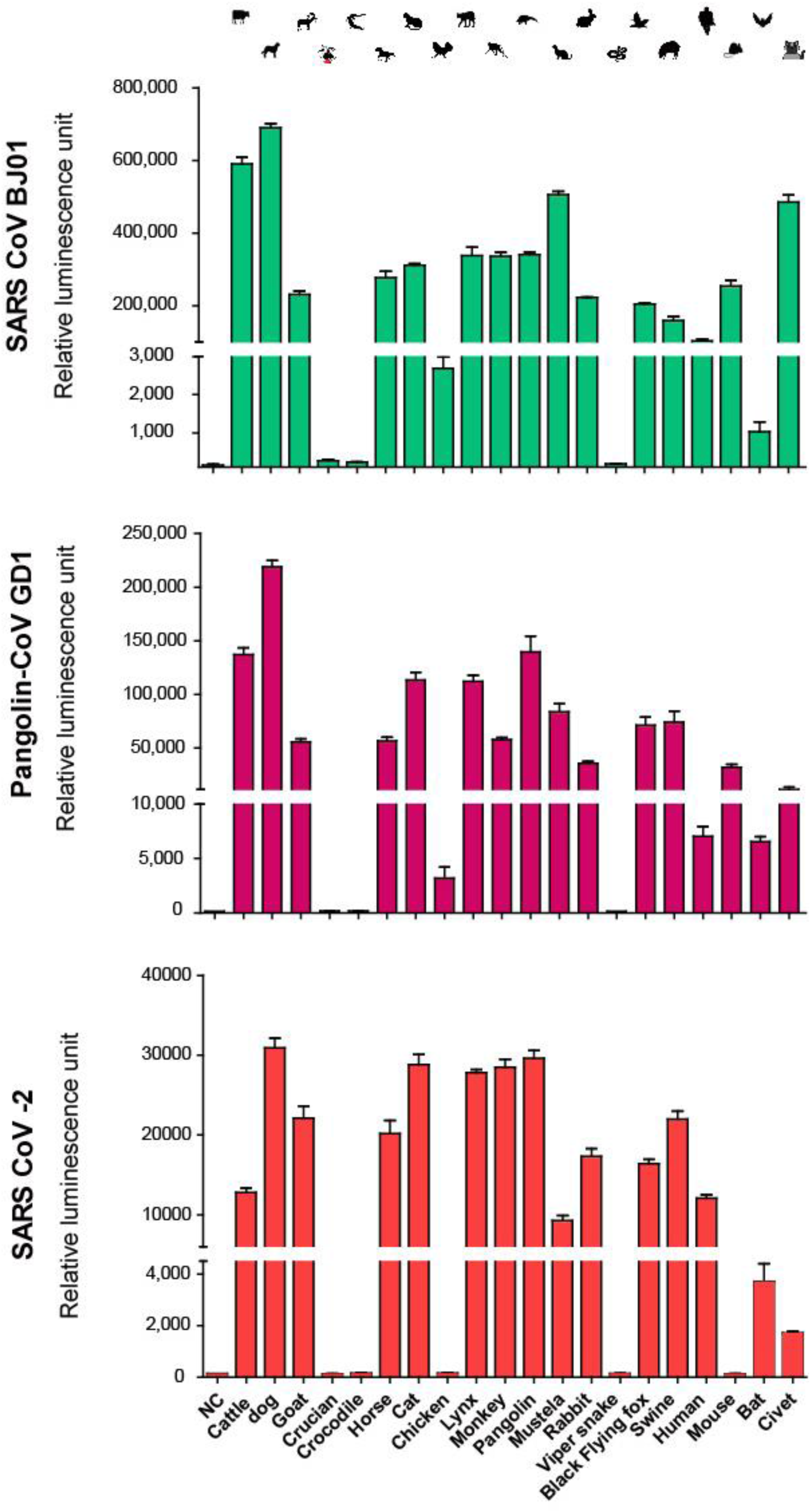
Entry efficiency of SARS-CoV-BJ01, SARS-CoV-2 and Pangolin CoV pseudoviruses into ACE2-expressing cells. HeLa cells expressing different ACE2 orthologs were infected by SARS-CoV, SARS-CoV-2 or Pangolin CoV pseudoviruses. At 48 h post infection, pseudovirus entry efficiency was determined by measuring luciferase activity in cell lysates. The results were presented as the mean relative luminescence units and the error bars indicated the standard deviations (n = 9).

### Phylogenetic analysis of ACE2s and key amino acids for SARS-CoV-2 utilization

To evaluate the phylogenetic relationship of the 20 ACE2s assessed above, we built a phylogenetic tree based on the aa sequences of all the ACE2s (**Figure 3A**). In the tree, we observed the obvious branches of mammalian, bird, reptilia and fish ACE2s. However, with the mammalian ACE2s, there were no significant branches corresponding to the ACE2 utilization by SARS-CoV-BJ01 or SARS-CoV-2 pseudoviruses, indicating that the whole sequence analysis could hardly reveal the key factors underlying the receptor utilization by the viruses.

**Figure 3.**
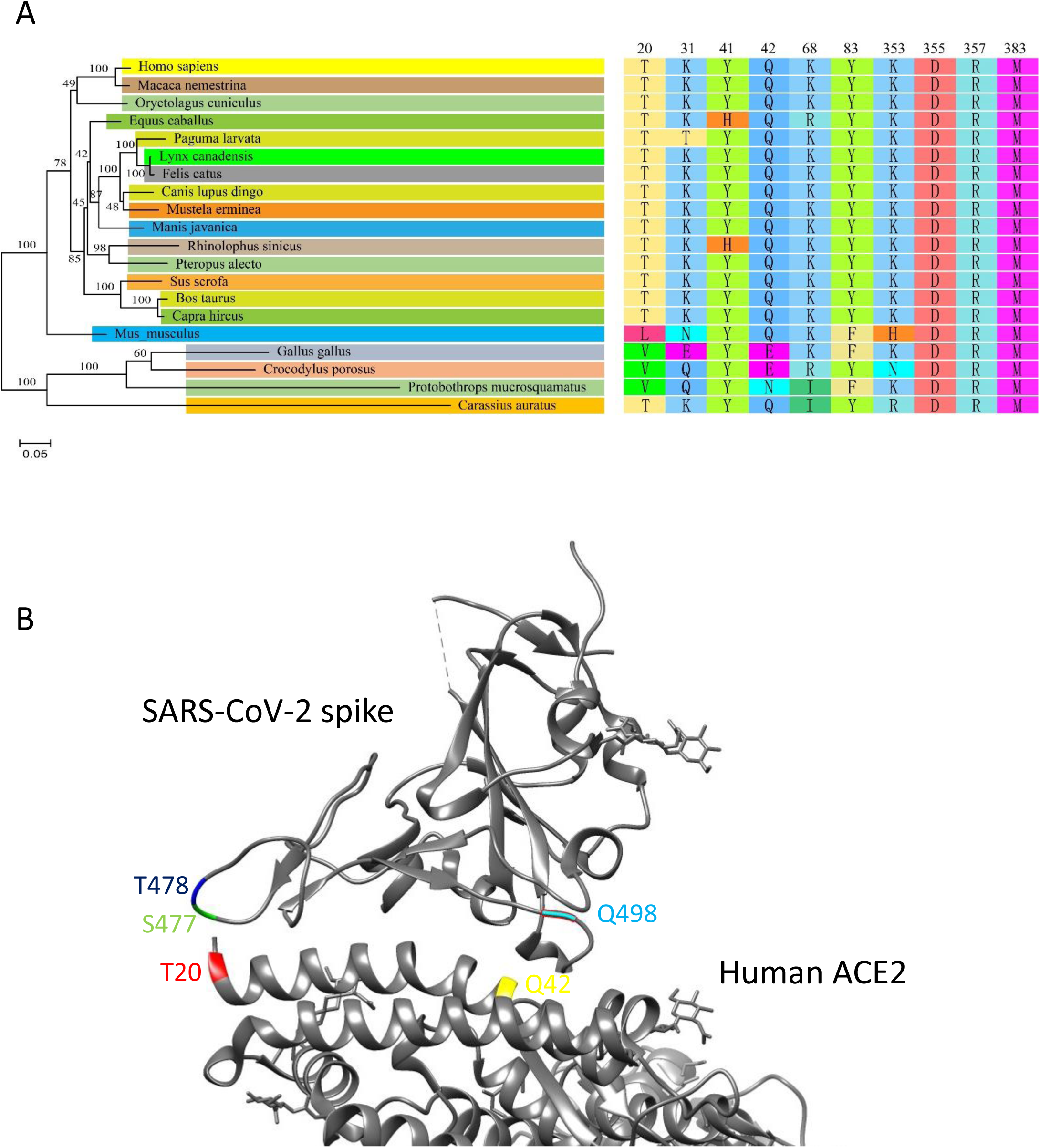
Phylogenetic analysis of the 20 ACE2 orthologs and the key amino acid residues for SARS-CoV-2 utilization. (A) The phylogenetic tree was constructed on the whole aa sequences of ACE2s using NJ method by MEGA7 with 1,000 bootstrap replicates (left panel) and the amino acids on the 9 critical sites predicted previously were listed (right panel). (B) The structure of the complex of SARS-CoV-2 spike and human ACE2 was adapted from Protein Data Bank (PDB ID: 6VW1). S477, T478 and Q498 of SARS-CoV-2 spike were labelled in green, blue and cyan, respectively. T20 and Q42 were labelled in red and yellow, respectively.

In our previous study, we predicted 9 key aa sites on human ACE2 potentially critical for the receptor utilization, including T20, K31, Y41, K68, Y83, K353, D355, R357 and M383, based on the SARS-CoV-2 utilization of human, bat, civet, swine and mouse ACE2s (18). Here we tested 15 more species, and then we checked the 9 sites of the 20 ACE2 orthologs to validate the role of the nine sites in SARS-CoV-2 utilization. As shown in **Figure 3A**, Y/H41, D355, R357 and R383 were conserved in all ACE2s, indicating that these sites were not determining the receptor utilization by SARS-CoV-2. Similarly, K68 was conserved in both mouse and chicken ACE2s that could not be used by SARS-CoV-2 and was probably not important for SARS-CoV-2 utilization either. Q42 was conserved in mouse ACE2 but was substituted by E42 in chicken ACE2 whose role in SARS-CoV-2 might be complicated. On the contrary, T20, K31 and Y83 in usable ACE2s were distinct from the corresponding aa in unusable ACE2s, indicating their cirtical role in determining SARS-CoV-2 utilization.

## Discussion

In history, most pandemic viruses originated from wild reservoir animals and transmitted directly or indirectly to humans, such as Ebola virus from bats (19) and human immunodeficiency viruses from chimpanzees (20). The reason for this phenomena is probably that native human viruses have undergone long-term coevolution with humans and are unlikely to cause severe diseases to their hosts after multiple natural selections. Therefore, interspecies transmission is usually a prerequisite for a virus to cause a pandemic. A broad host range is supposed to lead to effective interspecies transmission of virus and such virus is more likely to cause a pandemic. The COVID-19 pandemic caused by SARS-CoV-2 is the most severe worldwide pandemic in the recent years surpassing the SARS pandemic in 2003, so it is likely to speculate that SARS-CoV-2 has a broad host range. Surprisingly, our cell-entry result showed that SARS-CoV-2 had a smaller range of ACE2 utilization than SARS-CoV. SARS-CoV-2 could not utilize mouse or chicken ACE2 which could be used by SARS-CoV, indicating a narrower host range of SARS-CoV-2, especially in murine and birds. The reason is probably that the host range of SARS-CoV-2 is broad enough to support its transmission from bats to humans, and lack of infection to some kinds of animals does not affect such transmission due to redundant routes. Thus, it is not suggested to over-interpret the determination of the host range on the possibility of a virus to cause pandemics, especially for the viruses with broad host ranges.

Notably, SARS-CoV-2 has a better utilization of bat ACE2 than SARS-CoV. Though SARS-CoV originate from bat-SARSr-CoV, the utilization of bat ACE2 by SARS-CoV is quite limited which is supported by the previous reports (10, 21). According our current results, SARS-CoV-2 utilizes Chinese horseshoe bat ACE2 much better than SARS-CoV, indicating a higher homology between SARS-CoV-2 and its ancestor. This speculation is supported by the phylogenetic analysis of viral genomes in the previous study (1).

Our cell-entry result showed that SARS-CoV and SARS-CoV-2 could use a wide variety of mammalian ACE2s, which is further supported by the reports about the susceptibility of various mammals to SARS-CoV-2 infection (22). However, the utilization of fish and reptilian ACE2s was quite poor for both SARS-CoV and SARS-CoV-2. This can be explained by the remote phylogenetic relationship between fish/reptilian ACE2s and mammalian ACE2s. By comparison, bird ACE2s have closer phylogenetic relationship with mammalian ACE2s, and thus, bird ACE2s could be used by some but not all SARSr-CoVs, such as SARS-CoV. This indicates that SARSr-CoVs are more likely to be transmitted by mammals and birds but not fish and reptiles, and more attention should be paid to domestic mammals and birds to prevent CoV pandemic.

Our previous study predicted 9 critical aa residues on ACE2 orthologs based on the aa sequence comparison of Chinese horseshoe bat, civet, swine and mouse ACE2s, since these were the only ACE2s tested by SARS-CoV-2 utilization at that time (18). In this study, we have tested 16 more ACE2s which could help to confirm or exclude critical aa residues found previously. K31 and Y83 in human ACE2 were confirmed by this study as critical aa residues for SARS-CoV-2 utilization. Substitutions of K31D and Y83F were reported to abolish or strongly inhibit SARS-CoV binding (23, 24). Here we have found more K31 substitutions abolishing SARS-CoV-2 utilization that chicken ACE2 with K31E, crocodile and viper snake ACE2s with K31Q and mouse ACE2 with K31N could not be utilized by SARS-CoV-2, matching our previous prediction. Considering both chicken and mouse ACE2s could be utilized by SARS-CoV, K31 may be a key aa residue leading to the different receptor utilization of SARS-CoV and SARS-CoV-2. Also, chicken, mouse and viper snake ACE2s with Y83F could not be utilized by SARS-CoV-2, consistent with our previous prediction and other former reports. On the contrary, though Y41A was reported to abolish SARS-CoV binding (23), horse ACE2 with Y41H could be used by SARS-CoV-2 well, indicating that Y41H barely affect SARS-CoV-2 utilization. However, Chinese horseshoe bat ACE2 with Y41H showed a relatively lower utilization by SARS-CoV-2 and even lower utilization by SARS-CoV, indicating Y41H might decrease receptor utilization synergistically with only with other unknown substitutions, especially for SARS-CoV. Substitution of K68 was reported to slightly inhibit SARS-CoV binding (23). Nevertheless, horse ACE2 with K68R could be used by SARS-CoV-2 well, indicating that K68R might not affect SARS-CoV-2 utilization either. Crucian and viper snake ACE2 with K68I could not be used by SARS-CoV-2, but these ACE2s contains so many other substitutions that we may not attribute the abolishment of SARS-CoV-2 utilization to K68I.

Remarkably, our study revealed T20 of ACE2 as a potential key aa residue for SARS-CoV-2 utilization. T20 has not been reported to affect SARS-CoV utilization but mouse ACE2 with T20L and chicken ACE2 with T20V could not be used by SARS-CoV-2. T20 is located at the N terminus of most mature mammalian ACE2s (excluding the signal peptide). According to the structure of SARS-CoV-2 receptor-binding domain complexed with human ACE2, the N-terminal T20 of ACE2 is close to S477 and T478 of SARS-CoV-2 RBM (**Figure 3B**) (25). Both threonine and serine contain hydroxyl radicals that allow them to form hydrogen bonds with each other. Thus, T20 of human ACE2 is likely to bridge with S477 and T478 of SARS-CoV-2 spike via hydrogen bonds, which stabilizes the ACE2-spike binding. However, L20 on mouse ACE2 and V20 on chicken ACE2 are both aliphatic aa that cannot form hydrogen bonds to support the ACE2-spike binding, which may impair the utilization of these two ACE2s by SARS-CoV-2. In SARS-CoV spike, the two aa residues are substituted by G463 and K464. K464 can form hydrogen bonds with threonine and G463 is a non-polar aa that can interact with valine or leucine via hydrophobic bond together with the adjacent A461. Therefore, SARS-CoV spike can interact with various N-terminal aa of ACE2s, and this may be the reason why SARS-CoV can utilize mouse and chicken ACE2s while SARS-CoV-2 cannot. Pangolin CoV spike shares high similarity with SARS-CoV-2 which keeps S477 and T478. However, Pangolin CoV spike harbors alkaline H498 that can interact with acidic Q42 and E42 of mouse and chicken ACE2, respectively, via ionic affinity, which may complementally support the ACE2-spike binding and allow the utilization of mouse and chicken ACE2 by Pangolin CoV. On the contrary, SARS-CoV-2 spike substitutes H498 with acidic Q498, leading to ionic repulsion with Q42 and E42, blocking its binding with mouse and chicken ACE2s. These proposed interactions provide a possible explanation for the different utilization of ACE2s by SARS-CoV-2 and pangolin CoV. Nevertheless, all the speculation about the aa residues on ACE2 key for spike-ACE2 binding is based on the sequence analysis, and mutation validation is necessary to further verify the roles of those sites.

In summary, our current study has validated the ACE2 utilization by SARS-CoV-2 predicted in our previous analysis by pseudovirus cell-entry assay. The results showed less ACE2 utilization by SARS-CoV-2 compared to SARS-CoV and pangolin CoV, especially for murine and bird ACE2s, indicating narrower host range of SARS-CoV-2. Meanwhile, with these data, we screend 5 key aa residues of ACE2 for SARS-CoV-2 utilization. Especially, the N-terminal T20 and Q42 might be critical in determining the difference of ACE2 utilization by the three SARSr-CoV. Our findings deepen the understanding about the receptor utilization and the host range of SARS-CoV-2, providing useful information for tracing virus transmission routes and preventing pandemics caused by CoVs in the future.

## Methods and materials

### Cell lines and plasmids

HEK293T and HeLa cells were obtained from the American Tissue Culture Collection and cultured in Dulbecco’s modified Eagle medium (DMEM) (Gibco, USA) supplemented with 10% fetal bovine serum (FBS) in a humidified 5% CO2 incubator at 37°C.

Full-length SARS-CoV-2 spike (GenBank accession number MN908947.3), SARS-CoV BJ01 spike (GenBank accession number AY278488.2) and Pangolin CoV GD1 spike (GISAID accession number: EPI_ISL_410721) were all synthesized by Sangon and subcloned into the pcDNA3.1 vectors with a C-terminal HA tag. The cDNAs encoding different ACE2 proteins (Table S1) were synthesized by Sangon and subcloned into pcDNA3.1 vectors with a C-terminal His tag. All the plasmids were verified by Sanger sequencing.

### Western blot analysis

Lysates of cells or filtered supernatants containing pseudoviruses were separated by SDS–PAGE, followed by transfer to a nitrocellulose membrane (Millipore, USA). For detection of S protein, the membrane was incubated with anti-HA tag mouse monoclonal antibody (bimake, USA, 1:2000), and the bound antibodies were detected by Horseradish Peroxidase (HRP)-conjugated goat anti-mouse IgG (Abbkine, China, 1:5,000). For detection of HIV-1 p24 in supernatants, monoclonal antibody against HIV p24 (p24 MAb) was used as the primary antibody at a dilution of 1:8,000, followed by incubation with HRP-conjugated goat anti-mouse IgG at the same dilution. To detect the expression of 20 ACE2s in HeLa cells, Mouse anti-His tag monoclonal antibody (Bioworld, USA, 1:5000) was used as the first antibody, followed by incubation with HRP-conjugated goat anti-mouse IgG at the same dilution.

### Pseudovirus preparation and cell-entry assay

The pseudovirus cell-entry assay was performed as described previously (26). In brief, HEK293T cells were co-transfected with a luciferase-expressing HIV-1 plasmid (pNL4-3.Luc.R-E-) and a plasmid encoding HA-tagged SARS-CoV-BJ01 spike, SARS-CoV-2 spike or Pangolin CoV spike. The supernatant containing pseudoviruses was collected 48 h after transfection and the remaining cell pellet was lysed for Western blot detection of HA-tagged spike proteins. In cell-entry assay, pseudoviruses were incubated with recipient cells at 37 °C for 6 h, the medium was changed and cells were incubated for an additional 42 h. Cells were then washed with PBS buffer and lysed. Lysates were tested for luciferase activity (Promega, USA). Each infection experiment was carried out on for three times.

### Phylogenetic analysis

Multiple sequence alignment was performed for the whole aa sequences of ACEs using MAFFT with a local alignment strategy FFT-NS-2. The phylogenetic tree was constructed by MEGA7 using the neighbor-joining (NJ) method with 1,000 bootstrap replicates and visualized using FigTree.

## Acknowledgements

This work was jointly funded by the National Natural Science Foundation of China (grant number 32041001 and 81902070) and the Provincial Natural Science Foundation of Hunan Province (grant number 2019JJ20004, 2019JJ50035, and 2020SK3001).

## Author contributions

Conceptualization, Y.Q., Q.W., and X.G.; Lab Work, Y.Q., Q.W., J.L., and C.L.; Data Analysis, Y.Q., Q.W., and Z.Z.; Funding Acquisition, Y.Q. and X.G.; Writing - Original Draft, Y.Q.; Writing - Review & Editing, Y.Q. and X.G.

## Declaration of interest statement

The authors declare no competing interests.

## Notes

### Competing Interest Statement

The authors have declared no competing interest.

