## Supplementary material for "Receptor utilization of angiotensin converting enzyme 2 (ACE2) indicates a narrower host range of SARS-CoV-2 than that of SARS-CoV": Table S1

**Table S1**. **Information about the ACE2 orthologs used in this study**

| Species | Common Name | NCBI Accession |
|---|---|---|
| *Bos Taurus* | Cattle | NC_037357.1 |
| *Canis lupus dingo* | Dog | NW_020267596.1 |
| *Capra hircus* | Goat | NW_017189516.1 |
| *Carassius auratus* | Crucian | NC_039253.1 |
| *Crocodylus porosus* | Crocodile | NW_017728906.1 |
| *Equus caballus* | Horse | NC_009175.3 |
| *Felis catus* | Cat | NC_018741.3 |
| *Gallus gallus* | Chicken | NC_006088.5 |
| *Lynx canadensis* | Lynx | NC_044321.1 |
| *Macaca nemestrina* | Monkey | NW_012017243.1 |
| *Manis javanica* | Pangolin | NW_016527422.1 |
| *Mustela erminea* | Mustela | NC_045635.1 |
| *Oryctolagus cuniculus* | Rabbit | NC_013690.1 |
| *Protobothrops mucrosquamatus* | Viper snake | NW_015387079.1 |
| *Pteropus alecto* | Black flying fox | NW_006435837.1 |
| *Sus scrofa* | Swine | NC_010461.5 |
| *Homo sapiens* | Human | NM_021804.2 |
| *Mus musculus* | Mmouse | NM_001130513.1 |
| *Rhinolophus sinicus* | bat | KC881004.1 |
| *Paguma larvata* | civet | AY881174.1 |
